# Ouabain enhances cell-cell adhesion mediated by β_1_-subunits of the Na^+^,K^+^-ATPase in CHO fibroblasts

**DOI:** 10.1101/552497

**Authors:** Claudia Andrea Vilchis-Nestor, María Luisa Roldán, Teresita Padilla-Benavides, Liora Shoshani

## Abstract

Adhesion is an important characteristic of epithelial cells to provide a crucial barrier to pathogens and substances. In polarized epithelial cells, cell-adhesion depends on tight junctions, adherent junctions and the Na^+^,K^+^-ATPase. All these are located in the basolateral membrane of the cells. The hormone ouabain, a cardiotonic steroid, binds to the α subunit of the Na^+^,K^+^-ATPase, and inhibits the pump activity when used at above μM concentrations. At physiological nM concentrations, ouabain affects the adhesive properties of epithelial cells by inducing the expression of cell adhesion molecules through activation of signaling pathways associated to the α subunit. Our group showed that non-adherent CHO cells transfected with the canine β_1_ subunit become adhesive, and that homotypic interactions between β_1_ subunits of the Na^+^,K^+^-ATPase occur between neighboring epithelial cells. Therefore, in this study we investigated whether the adhesion between β_1_ subunits was also affected by ouabain. We used CHO fibroblasts stably expressing the β_1_ subunit of the Na^+^,K^+^-ATPase (CHO-β_1_) and studied the effect of ouabain on cell adhesion. Aggregation assays showed that ouabain increased the adhesion between CHO-β_1_ cells. Immunofluorescence and biotinylation assays showed that ouabain (50 nM) increases the expression of the β1 subunit of the Na^+^,K^+^-ATPase at the cell membrane. We also screened the effect of ouabain on activation of signaling pathways in CHO-β_1_ cells, and their effect on cell adhesion. We found that c-Src, is activated by ouabain and is therefore likely to regulate the adhesive properties of CHO-β_1_ cells. Collectively, our findings suggest that the β_1_ subunits adhesion is modulated by the levels of expression and activation of the Na^+^,K^+^-ATPase at the plasma membrane, which is regulated by ouabain.

## Introduction

The Na^+^,K^+^-ATPase or sodium pump is an ubiquitous plasma membrane transporter that creates the ionic gradients that drive the net movement of glucose, amino acids, and ions across cellular membranes [1]. The Na^+^,K^+^-ATPase belongs to the P-type ATPase family, whose members are characterized by the transitory formation of a phosphorylated enzyme intermediate [2]. The pump structurally consists of three subunits: a catalytic α subunit, an accessory β subunit and a regulatory γ subunit. The catalytic α subunit is constituted by 10 transmembrane domains (TM), and exchanges 3 Na^+^ ions from the cytosol for 2 K^+^ ions from the extracellular milieu using the energy released from ATP hydrolysis [3]. The β subunit is constituted by a single TM domain, and a long glycosylated extracellular domain. Its functions are discussed in detail below. The γ subunit is a small, single span TM protein belonging to the FXYD family, differentially expressed in tissues, which modulates the pump’s function [4, 5]. In mammals, there are four α subunit isoforms, three β subunit isoforms and seven FXYD members [6,7].

The β subunit of the sodium pump has different functions that depend on the type of isoform expressed (β_1,_ β_2_ or β_3_), and on the accompanying isoform of α subunit (α_1_-α_4_) [8]. The main and essential function is to chaperone for the α subunit [9] contributing to the assembly and arrival of the pump to the plasma membrane [10]. In addition, the β subunit undergoes conformational changes during the catalytic cycle [11]. Different β isoforms have been associated to different K^+^ affinities [12]. Furthermore, some studies suggest that the β_1_ subunit regulates cell polarity, cell motility, epithelial to mesenchymal transition, and oncogenic transformation [13–15]. In epithelia, the β_1_ isoform functions as a homophylic cell adhesion molecule [16,17]. Moreover, the β_2_ isoform is an <u>a</u>dhesion <u>m</u>olecule <u>o</u>n <u>g</u>lia (AMOG, [18]).

Emerging evidence showed that the Na^+^,K^+^-ATPase may have additional regulatory functions other than pumping ions across cell membranes. Ouabain and related cardiotonic steroids are highly specific Na^+^,K^+^-ATPase ligands that bind to all catalytic α isoforms [19–21]. Studies from various laboratories have documented an important signaling function of the Na^+^,K^+^-ATPase [8,22]. In epithelia, the sodium pump also acts as a membrane receptor that transduces signals in response to ouabain and related cardiotonic steroids. The binding of ouabain and cardiotonic steroids (at nM concentrations) to the sodium pump activates signaling pathways that resemble those triggered by hormone/receptor interactions, which regulate gene expression, membrane trafficking, cell adhesion, proliferation, and cell death [23–28]. Interestingly, nM concentrations of ouabain neither inhibit K^+^ pumping nor disturb the K^+^ balance of the cell [29]. Therefore it was proposed that low ouabain concentrations bind and activate a non-pumping population of the Na^+^,K^+^-ATPase [30, 31].

Ouabain was suggested to be a hormone when Hamlyn and Mathews demonstrated the presence of a substance in plasma similar to ouabain of vegetal origin [20,32]. Thereafter, it was demonstrated that endogenous ouabain is synthesized and secreted by the hypothalamus [33,34], and the adrenocortical gland [35–37]. The status of ouabain as a hormone was clearly recognized upon the discovery of increased concentrations in plasma during exercise, salty meals ingestion, and pathological conditions such as arterial hypertension and myocardial infarction [38–43]. However, its physiological role remained unknown. Work from our group has shown that ouabain binding to the Na^+^,K^+^-ATPase modulates epithelial cell adhesion and communication [29, 44,45].

Our laboratory has studied the role of the Na^+^,K^+^-ATPase β_1_ subunit in epithelia. We demonstrated that the β_1_-subunits of Na^+^,K^+^-ATPases on neighboring cells interact with each other in a species specific manner [46,16]. Numerous studies demonstrated that the intercellular homotypic interaction between β_1_ subunits of the Na^+^,K^+^-ATPase are important for the stability of adherent junctions (AJ) and integrity of the tight junctions (TJ) and GAP junctions [17, 45, 47]. Thus, β_1_-β_1_ interactions between epithelial cells are critical for the integrity of intercellular junctions. Since ouabain modulates different cell-attachment complexes, we wondered whether ouabain also regulates the β_1_-β_1_ mediated cell adhesion. In this work we used CHO fibroblasts, which lack the classical cell-cell adhesion complexes (TJs, AJ), and that stably express a canine β_1_ subunit in the plasma membrane. This model system targets efficiently the Na^+^,K^+^-ATPase to the membrane contributing to the cell-cell contact [46]. To determine whether β_1_-β_1_ interactions are modulated by ouabain, we investigated the effect of a physiological concentration (50 nM) of ouabain on the adhesion of CHO cells overexpressing the dog β_1_ subunit. This work shows that ouabain increases the amount of Na^+^,K^+^-ATPase at the cell membrane, rendering increased cell adhesion properties mediated by β_1_-β_1_ interactions. This effect is facilitated by the ouabain-dependent activation of kinases such as cSrc, ERK1/2 and AKT which contribute to enhance the adhesive properties of CHOβ_1_ cells.

## Results

### Cell-Cell adhesion of CHO fibroblasts expressing dog β_1_ subunit of Na^+^,K^+^-ATPase is mediated by β_1_ homotypic interactions in trans

Adherent CHO fibroblasts are well attached to the extracellular matrix and the substrate, but establish weak cell-cell contacts, which are easily disrupted by gentle shaking or pipetting [46, 52, 53]. We have shown that CHO fibroblasts transfected with the canine β_1_ subunit of the Na^+^,K^+^-ATPase (CHOβ_1_) form large cellular aggregates, due to an increase in cell-cell adhesion [46]. Moreover, we demonstrated that the epithelial β_1_ subunit of Na^+^,K^+^-ATPase is an adhesion molecule that mediates the interaction of sodium pumps on neighboring cells by establishing homotypic interactions [16]. Therefore, we hypothesized that the cell-cell adhesion observed in CHOβ_1_ cells is mediated by β_1-_β_1_ interactions. To address this question, we investigated the subcellular localization of the dog β_1_ subunit and the cell-cell adhesion properties of CHO cells. Wild type CHO cells do not express the β_1_ subunit of the sodium pump (Fig. 1A left panel). Confocal microscopy analyses showed that CHO cells transfected with the plasmid encoding for the canine β_1_ subunit, express the protein in the plasma membrane, resembling the distribution observed in epithelial MDCK cells (Fig. 1A Middle and right panels). Dispase assays showed that wild type CHO cells are unable to maintain cellular aggregates (Fig. 1B left panel). However, the cell-cell adhesion capability of CHO cells increased upon transfection of the canine β_1_ subunit, as these cells maintain larger cellular aggregates upon dispase disruption, similar to those of the CHO cells expressing the adhesion molecule E-cadherin (CHO E-cadh; Fig. 1B). On average, the aggregates formed by CHOβ_1_ cells are significantly larger (3 fold) than control wild type CHO cells; but similar in size to those observed in the CHO E-cadh cells (Fig. 1C).

**Figure 1.**
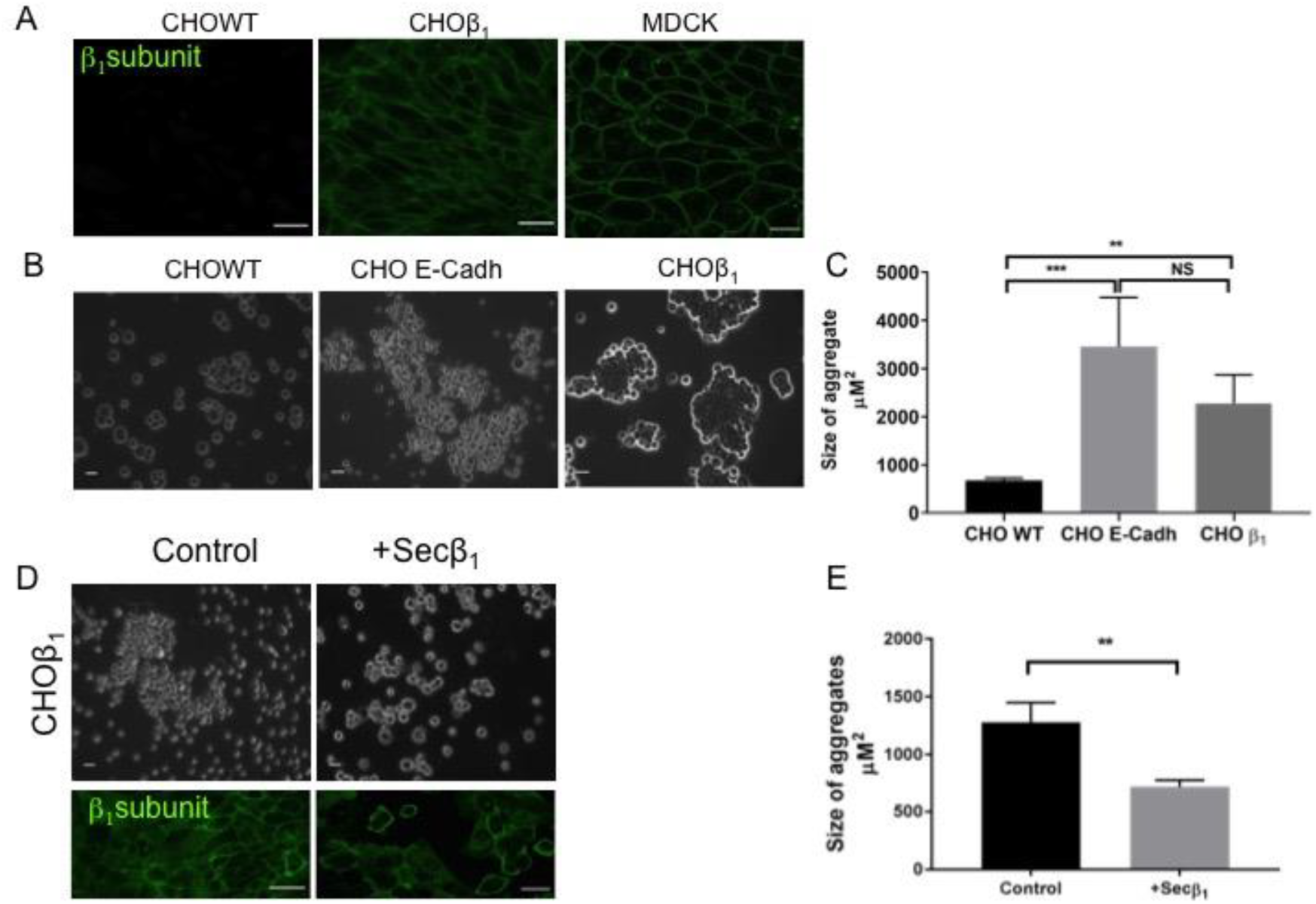
Cell-cell adhesion of CHO fibroblasts expressing the canine β_1_ subunit of Na^+^,K^+^-ATPase (CHOβ_1_) is mediated by β_1_-β_1_ interactions. (A) Representative confocal microscopy images of CHO wild type, CHOβ_1_ and MDCK cells immunostained against the dog β_1_ subunit. Scale bar=30 μm. (B) Representative light microscopy images of CHO wild type, CHOβ_1_ cells and CHO E-Cadherin after the dispase aggregation assay. Scale bar=20 μm. (C) Quantification of the measured size of the aggregates is depicted as the area of their horizontal projections. Values represent the mean from three independent biological replicates ± SE. Statistical analysis was Kruskal Wallis and Dunnet’s t-test for multiple comparisons; P<0.005. (D) Upper panels are representative phase-contrast micrographs of aggregation assay as in (B). Scale bar = 20 μm. Lower panels are representative confocal microscopy images of dog β_1_ subunit in CHOβ_1_ cells incubated for 24 h in the absence (left) or presence of Secβ_1_ (right). (E) Quantification of the mean size of cellular aggregates of CHOβ_1_ cells treated or not with Secβ_1_. Statistical analysis was t-student test of three independent biological experiments ± SD; P <0.005.

To confirm the hypothesis that the cell-cell adhesion observed in CHOβ_1_ cells is due to β_1-_β_1_ interactions, we tested whether the soluble domain of the β_1_ subunit would impair the formation of cellular aggregates in this cell line. We took advantage of a truncated version of the canine β_1_ subunit that only expresses the soluble extracellular C-terminal domain (Sec β_1_) [16, 51]. CHOβ_1_ cells were allowed to interact with supernatants obtained from CHO Secβ_1_ cells containing this protein, and the formation of cellular aggregates was analyzed by light microscopy. Figure 1D shows that the presence of the soluble domain of the dog β_1_ subunit (Secβ_1_) reduced the size of the CHOβ_1_ cellular aggregates. Statistical analyses confirmed that the aggregates formed by CHOβ_1_ cells were significantly smaller (~50%) than control cells (Fig. 1E). Interestingly, confocal microscopy analyses showed that CHOβ_1_ cells pre-incubated for 24 h with Secβ_1_ presented a proliferation defect compared to control cells (Fig. 1D, lower panel). These results confirmed that cell-cell adhesion dependent in the Na^+^,K^+^-ATPase is at least partially due to an interaction between β_1_ subunits, and further showed that the cell culture model based on CHOβ1 cells is suitable for studying β_1_-β_1_ interactions.

### Ouabain increases cell-cell adhesion of CHOβ1 cells

Physiological concentrations of ouabain modulate cell-cell interactions [26,29]. Therefore, we hypothesized that ouabain may also modulate the cell-cell interactions that are mediated by the β_1_ subunits of the sodium pump. To test this hypothesis, we used the dispase adhesion assay, to further investigate the adhesive properties of CHOβ_1_ cells in absence or presence of ouabain. Figure 2 shows a specific and significant increase of the size of CHOβ1 cells aggregates upon treatment with ouabain at physiological concentration (50 nM; compare Fig. 2A upper and lower panels). Importantly, the inhibitory ouabain concentration (100 μM) prevented the cell adhesion phenotype observed in CHOβ_1_ cells, to a level similar to wild type cells. Therefore, we concluded that physiological concentrations of ouabain increase cell adhesion mediated by β_1_-β_1_ interactions of CHOβ_1_ cells.

**Figure 2.**
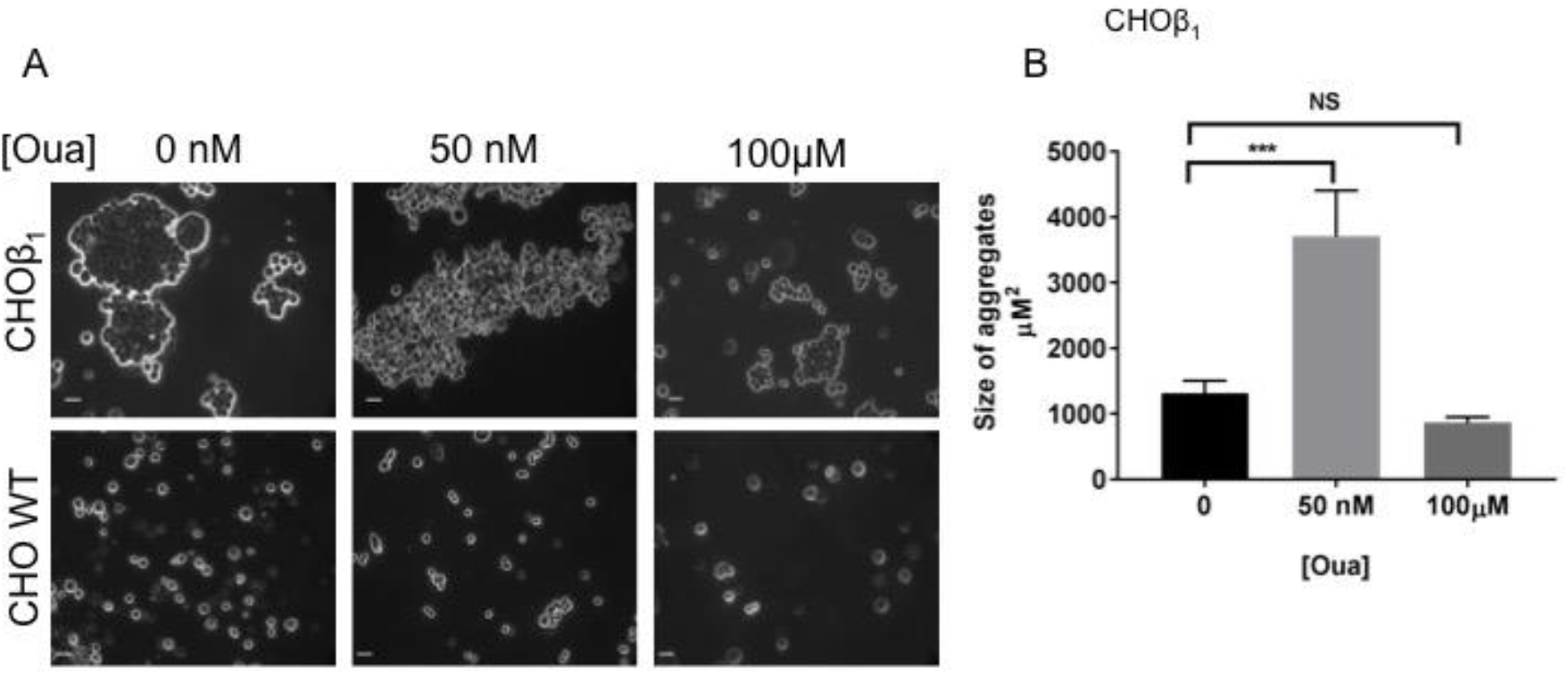
Physiological concentrations of ouabain increase cell-cell adhesion of CHOβ_1_ cells. (A) Representative microscopy images of cell aggregation of CHOβ_1_ cells (upper panels) and CHO WT cells (lower panels) treated or not with ouabain (50 nM, 100μM). (B) Quantification of the mean size of cellular aggregates of CHOβ_1_ treated with 0, 50nM and 100μM ouabain of three independent biological replicates is depicted. Statistical analysis was ANOVA one way and Dunn’s tests for multiple comparison, bars represent ± SE; *** P<0.0001.

### The interactions of β_1_-β_1_ subunits are stable in vitro independently of ouabain treatment

The effect of ouabain on cell-cell interactions mediated by β_1_ subunits could be explained if this hormone induces a conformational change in β_1_ subunits resulting in a more adhesive molecule. Accordingly, ouabain binding to its receptor, the α subunit of the pump, should be sufficient for inducing the same effect on β_1_-β_1_ interactions *in vitro*, out of the cellular context. Therefore, we studied whether ouabain directly affects β_1_-β_1_ subunits interaction in a pool down assay. In this case, we used the canine β_1_ tagged with a hexa-histidine repeat (CHOβ_1_ His_6_) immobilized on Ni^+^-NTA as the bait. The prey was obtained from total cellular extract of CHOβ_1_ cells tagged with the yellow fluorescent protein (CHOβ-YFP). CHO wild type cells (CHO WT) were used as negative control. Cellular extracts (bait and prey) were allowed to interact in absence or presence of ouabain, and the formation of complexes interacting β_1_ subunit were analyzed by Western blot. Figure 3 shows that the immobilized CHOβ_1_ His_6_ was capable to interact with the recombinant β_1_ YFP obtained from cellular extracts. Importantly, the interaction *in vitro* was maintained even in the presence of ouabain (Fig. 3A). Statistical analyses demonstrated that ouabain treatment did not affect significantly the amount of interacting proteins (Fig. 3B). All eluted fractions contain α subunit, which means that the β_1_His_6_ is assembled with the α subunit on the Ni^+^-NTA. The data suggest that the effect of ouabain on cell-adhesion does not occur directly on β_1_ subunits, and it is likely dependent on additional cellular components.

**Figure 3.**
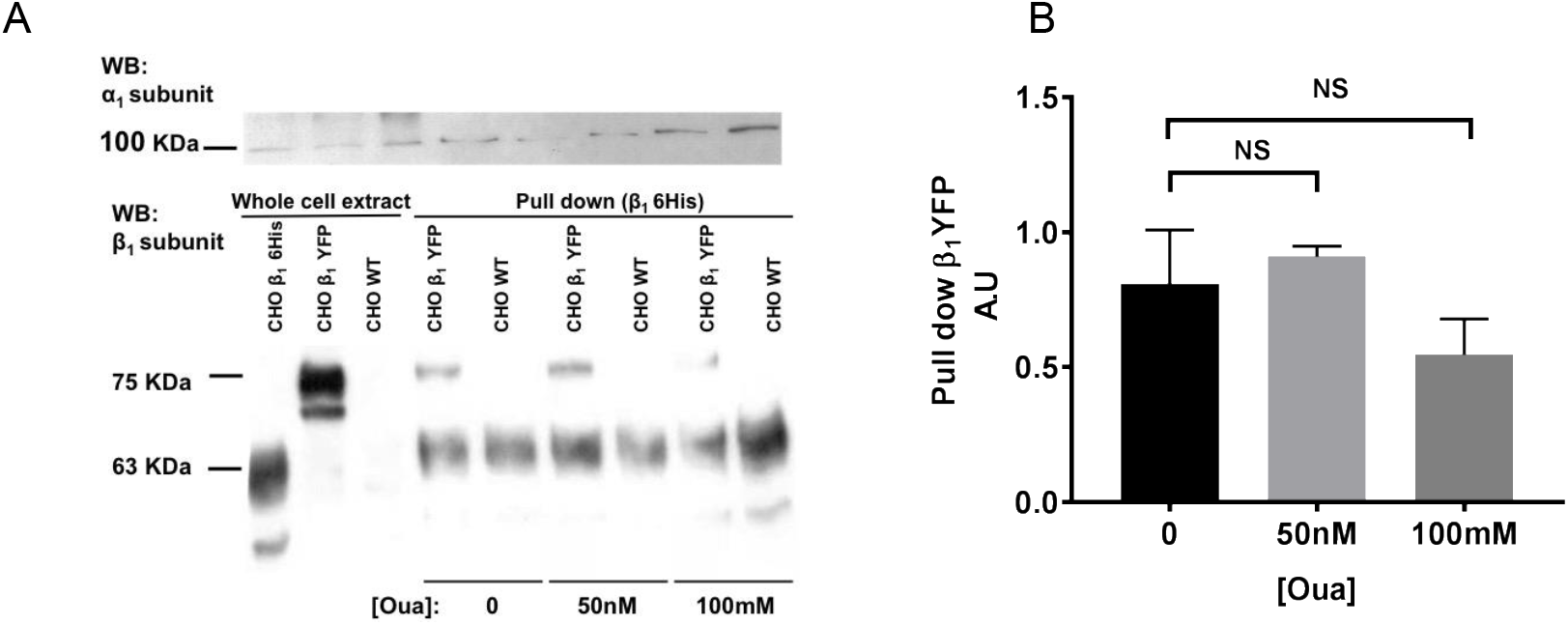
An indirect interaction between ouabain and the β_1_ subunit of Na^+^,K^+^-ATPase is necessary for the adhesive phenotype triggered by this hormone. (A) Western blot analysis of β_1_ and α_1_ subunits of the Na^+^,K^+^-ATPase: cell lysate of CHOβ_1_-6His, CHOβ1-YFP and CHO-WT cells (three left lanes); Pull Down assay between CHOβ_1_-6His as a bait and CHOβ_1_-YFP or CHO-WT as a prey without and with 50 nM and 100 mM Ouabain (six right lanes). The 63 KDa band corresponds to the β_1_-6His, and the higher-molecular-weight band corresponds to the β_1_-YFP (75KDa) both recognized by the same anti-β_1_ subunit antibody. (B) Densitometric quantification of the results presented in A. In Pulled down fractions, density of the β_1_-YFP bands was normalized with the β_1_6His band. Data represent the mean of three independent replicates. Statistical analysis was Kruskal Wallis and Dunn’s test for multiple comparisons. Error bars, ± SE; NS, P>0.05.

### Ouabain increases the expression and localization of the sodium pump at the plasma membrane

It has been shown that treatment with cardiotonic steroids (ouabain and 21-benzylidene digoxin) increases the expression level of the α subunit of sodium pump in the typical pig kidney tubular epithelium cell line LLC-PK1 and MDCK [54, 55]. Therefore, we wondered whether the cell adhesion effect of ouabain induced in CHOβ_1_ cells was due to increase in the expression of the sodium pump at the plasma membrane. As expected, treatment of MDCK cells with ouabain at physiological concentrations (10-50 nM) increased the expression of the β_1_ subunit at the plasma membrane (Fig. 4A), which is in agreement with the observations of Rocha, and coworkers (2014). On the other hand, treatment with µM concentrations of ouabain leads to detachment of the MDCK cells (Fig. 4A; [56]). Considering these phenotypes, we evaluated the effect of ouabain on the localization of the β_1_ subunit in the monolayer of CHOβ_1_ cells. Confocal microscopy analyses showed that in our model system, the β_1_ subunit presents a similar distribution to epithelial cells upon ouabain treatments (Fig. 4B). The membranal localization of the sodium pump was disrupted at higher concentrations of ouabain (µM) (Fig. 4B). Importantly, the fluorescence intensity of the β_1_ subunit at the plasma membrane only increased significantly upon treatment with 50 nM ouabain (Fig. 4C).

**Figure 4.**
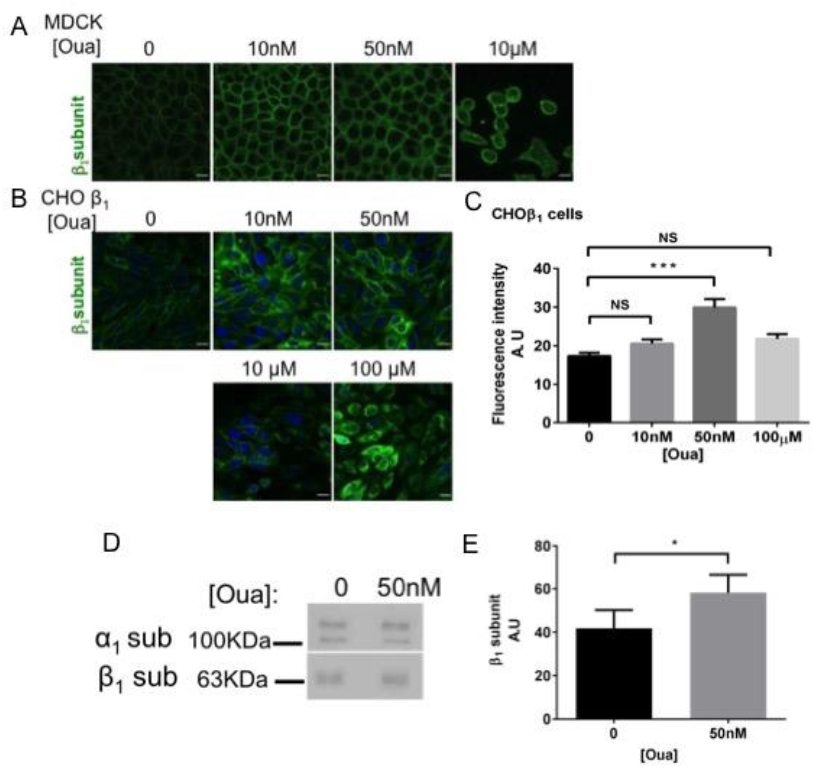
Ouabain increases the expression of the Na^+^,K^+^-ATPase at the plasma membrane of CHO fibroblasts transfected with the canine β_1_ subunit. (A) Representative confocal microscopy images of the dog β_1_ subunit in MDCK cells incubated with 10, 50 nM and 10 μM of Ouabain for 24 h. (**B**) Representative confocal microscopy images of the dog β_1_ subunit in CHOβ_1_ cells incubated for 24 h with 10, 50 nM, 10, and 100 μM of ouabain. (**C**) Quantification of the fluorescence intensity observed at the plasma membrane of CHOβ_1_ cells (10 cells per field of three independent biological replicates). Statistical analyses were ANOVA one way test, and Dunn’s test for multiple comparisons. The data represent the mean ± SE; ***P<0.0001, NS. (**D**) Representative Western blot analysis of β1 and α1 subunits of the Na^+^,K^+^-ATPase of CHOβ_1_ cells biotinylated after treatment with 50 nM ouabain for 24 h. (E) Densitometry quantification of biotinylated β_1_ subunit after treatment with ouabain. The data represent the mean of three independent biological replicates ± SD, Statistical analyses was t-student test; * P <0.05.

To confirm that physiological concentrations of ouabain increase the sodium pump expression at the plasma membrane, we used a surface biotinylation assay. Figure 4D shows a representative Western blot analysis of the β_1_ subunit located in the plasma membrane of CHOβ_1_ cells treated with 50 nM ouabain. Statistical analyses of three independent biological experiments showed a small but significant increase in the membranal expression of the β_1_ subunit in CHOβ_1_ cells (Fig. 4D). Our results suggest that the observed increase in cell adhesion mediated by β_1_ subunits is a result of ouabain binding to the α subunit, which may lead to the activation of signaling pathways that promote the expression and delivery of the pump to the plasma membrane.

### The β_1_-β_1_ adhesion induced by ouabain depends on the activation of cSrc and AKT signaling pathway

Previous studies demonstrated that binding of cardiotonic steroids such as ouabain to the sodium pump stimulates multiple kinase cascades [57]. Therefore, the effect of ouabain on the β_1_-β_1_ interaction could be explained by the potential activation of some components from the signaling machinery that is known to modulate cell junctions such as TJs, AJs and GAP junctions [29,58]. We first screened for signaling proteins of the family of tyrosine kinases that were differentially activated compering between CHOβ_1_ and CHO wild type cells. Overall, CHOβ_1_ cells had more active signaling proteins than wild type cells, indicating that the increased expression of Na^+^,K^+^-ATPase in CHO fibroblast per se induces the activation of signaling cascades that are usually of low activity (Figure 1). Then, we treated CHOβ_1_ cells with ouabain and identified various kinases that were altered when treating the cells with 50 nM Ouabain (Figure S2). Of all modified pathways detected, we focused on the potential mechanisms by which cSrc, AKT, and ERK1/2 contribute to the interaction between β_1_ subunits. Therefore, we investigated whether the inhibition of those pathways would impair the effect of ouabain on cell adhesion mediated by β_1_ subunits. The selected phosphorylation inhibitors were Dasatinib for cSrc, Perifosine for AKT, and U01266 for ERK1/2. From the three kinases selected, phosphorylated cSrc increased upon treatment with ouabain compared to control cells (Fig. 5A). Importantly, incubation of the CHOβ_1_ cells with all the inhibitors was effective and prevented the corresponding kinase phosphorylation (Fig. 5A-C).

**Figure 5.**
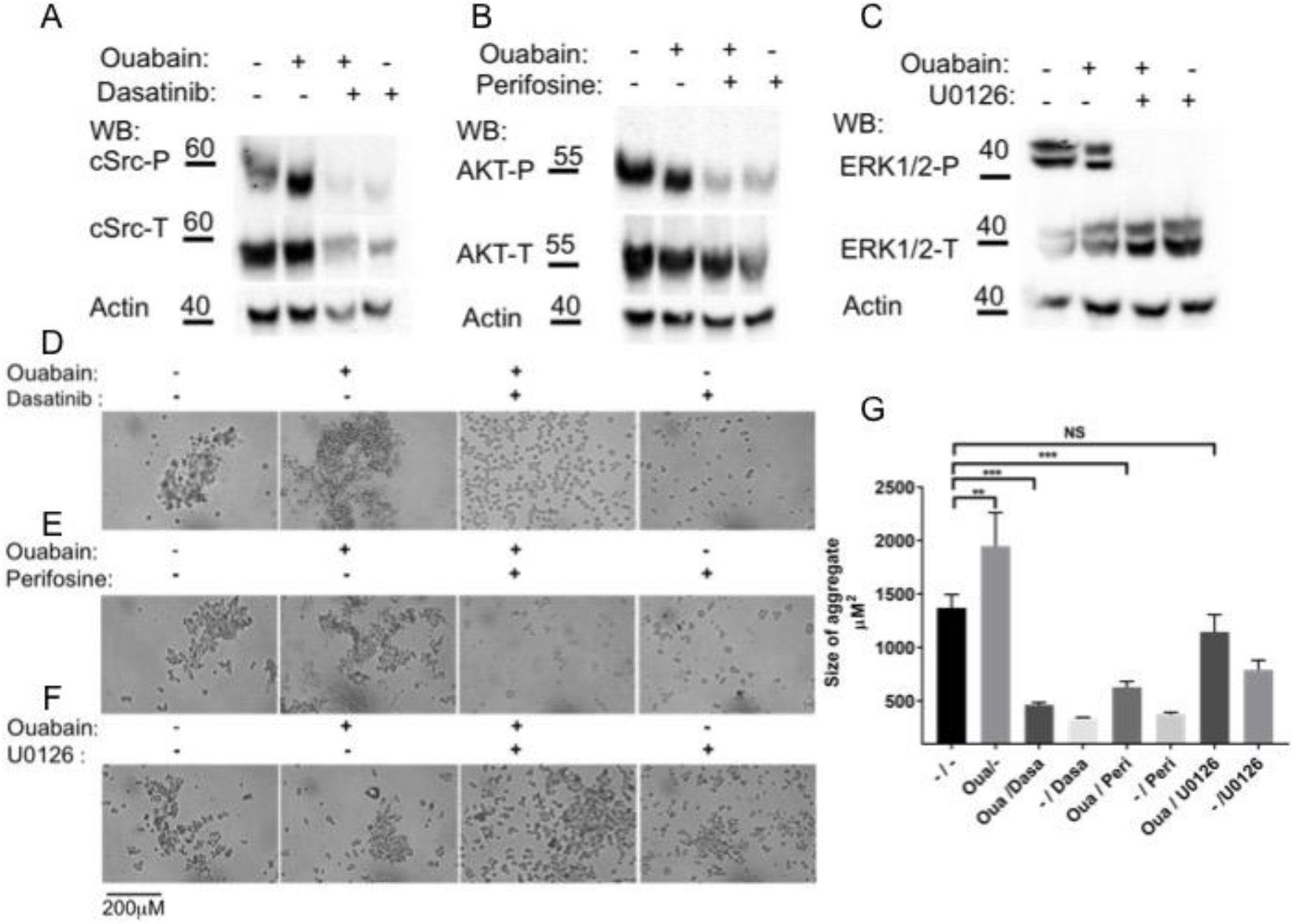
Src and AKT signaling pathways are involved in ouabain modulation of cell-cell Adhesion of CHOβ_1_ fibroblasts. Representative Western blot analysis of Src-P (A), AKT-P (B) and ERK1/2-P (C) of CHOβ_1_ cells treated without and with 50 nM ouabain, and in the presence or absence of the specific kinase inhibitor: Src (Desatinib, 100 nM); AKT (Perifosine, 10 μM); and EKR1/2 (U0126, 10 μM) (n=3). The inhibitors were added 1 h prior incubation with ouabain for 60 min. Representative images taken for Dispase aggregation assay of CHOβ_1_ cells treated with 50nM ouabain and the specific inhibitors for Src, Desatinib (D); AKT, Perifosine (E) and EKR1/2, U0126 (F). (G) Quantification of the size of cellular aggregates. The data represent the mean of three independent biological replicates ± SE. Statistical analysis was Kruskal Wallis and Dunn’s test for multiple comparisons. *** P<0.0001, ** P<0.001, NS P>0.05.

To test the effect of kinase inhibition in the adhesive properties of CHOβ_1_ cells dependent on ouabain (50 nM), we analyzed the size of the cellular aggregates. Representative light microscopy imaging showed that inhibition of cSrc (Fig. 5D), and AKT (Fig. 5E) impaired the formation of the ouabain-induced aggregates. No changes on the size of the cellular aggregates of CHOβ_1_ cells that were inhibited for ERK1/2 were observed (Fig. 5F). Consistently, statistical analyses showed significant differences in the size of the aggregates of CHOβ_1_ cells treated with either perifosine or dasatinib; no significant differences were observed for CHOβ_1_ cells treated with UO126 when compared to control cells (Fig. 5G). These data suggest that ouabain binding to the sodium pump stimulates at least cSrc kinase and AKT signaling pathways, which may be involved in the regulation of β_1_-β_1_ mediated cell adhesion.

### pNaKtide reduces cell adhesion effect induced by ouabain

The Na^+^,K^+^-ATPase interacts with cSrc kinase and forms a complex that serves as a receptor for ouabain to stimulate various protein kinase cascades [30]. Specifically, ouabain binding to the Na^+^,K^+^-ATPase disrupts this interaction and results in the assembly and activation of different signaling pathways [59]. Our results thus far have shown that the adhesive properties of β_1_ subunits require the cellular context to occur, since no effect of ouabain was observed *in vitro* (Fig. 3). Moreover, the increased phosphorylation of cSrc in CHOβ_1_ cells treated with ouabain (Fig. 5A), was reversed by the cSrc kinase inhibitor, and prevented formation of cellular aggregates (Fig. 5D). Therefore, we hypothesized that the β_1_ subunit adhesive properties dependent in ouabain, could be partially due to the activation of Na^+^,K^+^-ATPase/cSrc complex. To test this hypothesis we took advantage of the pNaKtide, which is a peptide that antagonizes the effect of ouabain on cSrc [60], and assayed for cell-adhesion in the presence of ouabain and with or without the pNaKtide. Figure 6A shows a representative Western blot of the ouabain-dependent increase in the phosphorylation of cSrc (~70% higher than control cells). In the presence of 1 μM of the pNaKtide, the effect on cSrc phosphorylation was abolished, and remained at similar levels than control cells. No changes were detected in the expression of the α subunit of the sodium pump. Finally, we analyzed the effect of the antagonist in the formation of ouabain-dependent aggregates of CHOβ_1_ cells. Representative light microscopy images showed that the size of the aggregates is smaller compared to those of ouabain stimulated cells (Fig. 6B). Statistical analyses showed that in the presence of the pNaKtide the size of ouabain-stimulated aggregates is similar in size than those of control cells (Fig. 6C). These results support our hypothesis that the adhesion effect of β_1_ subunits stimulated by ouabain is partially regulated by cSrc activation.

**Figure 6.**
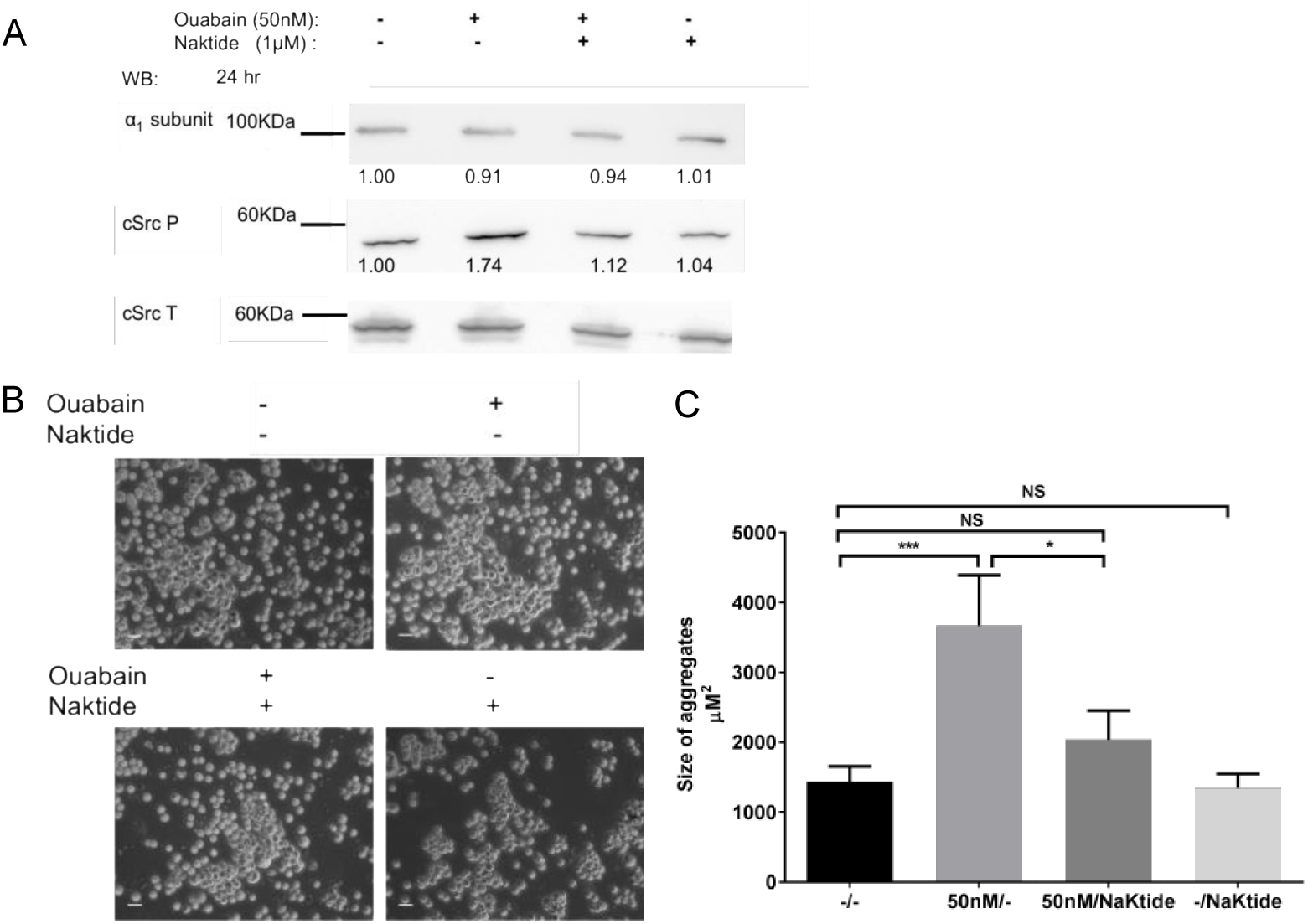
pNaKtide inhibits Ouabain-activated cSrc and ouabain induced β_1_-mediated cell adhesion in CHOβ_1_ cells. (A) Representative Western blot of cSrc activation (cSrcP) and total cSrc (cSrcT), and the α_1_ subunits of CHOβ_1_ cells treated with 1 mM pNaKtide for 24 h (n=3). (B) Representative microscopy images of the dispase assay of CHOβ_1_ cells treated with pNaKtide and ouabain for 24 h; scale bar = 20 μm. (C) Quantification of the size of cellular aggregates. The data represent the mean of three independent biological replicates ± SE. Statistical analyses was Kruskal Wallis and Dunn’s test for multiple comparisons. *** P<0.0001; * P<0.05; NS P>0.05.

## Discussion

As all mammalian cells, CHO fibroblasts express the necessary amount of sodium pumps at their plasma membrane for maintaining a suitable ionic balance that would permit their viability. In our overexpression system, the assembled α_1_β_1_ dimer in CHOβ_1_ cells is targeted to the plasma membrane and concentrated at cell-cell contacts adopting an epithelial-like phenotype and maintain both cell-cell and cell-substrate adhesion [46]. This phenotype makes the CHOβ_1_ overexpression system an ideal model to investigate cell adhesion. Using cellular and biochemical strategies our group demonstrated that β_1_ subunits of neighboring cells can interact directly in a species-specific manner [16]. In the present work we show that increasing the density of the β_1_ subunit of Na^+^-pump in cell contacts of CHO fibroblasts is sufficient for displaying a cell-adhesive phenotype. These adhesive properties are mediated by β_1_ interactions which are regulated by signaling cascades conserved in establishment and maintenance of classical epithelial adhesion complexes [26,29,45].

Early works from Nelson’s group on epithelial cell polarity have shown that expression of E-cadherin in L-fibroblasts, which lack surface polarity resulted in membrane domains with apical and basolateral identities [61]. Remarkably, the endogenous Na^+^-pumps in those experiments were recruited to cell-cell contacts. Our results show that CHO fibroblasts transfected with E-cadherin, a Ca^2+^-dependent cell adhesion molecule, or with the β_1_ subunit, a Ca^2+^-independent adhesion molecule, display similar aggregation properties. To validate that cell-cell adhesion of CHOβ_1_ cells was in fact due to β_1_-β_1_ interactions, cellular aggregation was challenged with Secβ_1_, the soluble extra-cellular domain of the canine β_1_ subunit. As expected, this truncated version of the β_1_ prevented the ouabain induced cell-cell adhesion properties of CHOβ_1_ cells. These results strongly suggest that secβ_1_ competes for cell-cell adhesion mediated by β_1_ subunits on neighboring cells. And that ouabain-induced cell-cell adhesion is indeed through β_1_-β_1_ interactions. Remarkably, Secβ_1_ blocks cell-cell aggregation between CHOβ_1_ cells only when it is added short time after seeding, before the formation of a confluent monolayer. This observation indicates that Secβ_1_ is not able to compete with already formed β_1_-β_1_ interactions but is able to interact with non-occupied β_1_ subunit and prevent cell-cell adhesion mediated by β_1_ subunits. Moreover, CHO β_1_ cells pre-incubated with secβ_1_ do not reach confluence and are apparently not proliferating. The cells form small patches and many single cells are observed. Usually, contact naïve epithelial cells do not express the Na^+^-pump at the plasma membrane [62]. Importantly, MDCK cells co-cultured with CHO wild type cells do not express the Na^+^,K^+^-ATPase at the heterotypic contacting membrane [46]. Nevertheless, some of the CHOβ_1_ single cells showed a highly fluorescent signal at the plasma membrane, which corresponded to the β_1_ subunit. Therefore, it is plausible that these contact naïve CHOβ_1_ cells, are actually surrounded by soluble Secβ_1_ molecules associated with the membrane-bound β_1_ subunit and that it is mimicking cell-cell contacts. As such, we hypothesize that the apparent proliferation defect is due to a contact inhibition effect induced by Secβ_1_. These data open a new paradigm on the modulation of cell proliferation dependent on the sodium pump. Altogether, these results confirm that cell-cell adhesion of CHO fibroblasts that overexpress Na^+^,K^+^-ATPase is mainly based on β_1_-β_1_ interactions between neighboring cells.

Ouabain binding to α subunit of the epithelial Na^+^,K^+^-ATPase has a dual effect on the pump, high concentrations of ouabain (>300 nM) triggers pump inhibition and cell detachment [44,56]. Low concentrations of ouabain increases sodium pump activity [63,64] and stimulate signaling routs in a cellular context dependent manner [65]. In epithelial MDCK cells, low ouabain concentrations (10-100 nM) modulate cell-cell contacts such as TJs, AJs and GAP Junctions [26, 29, 44, 45]. We analyzed the effect of ouabain on β_1_-β_1_ mediated cell adhesion. Our results show that CHOβ_1_ cells incubated with low concentration of ouabain (50 nM) make bigger aggregates than the non-treated ones, while incubation with high concentration of ouabain (100 µM) does not enhance cell-cell adhesion. Thus we conclude that physiological concentrations of ouabain induce an increase in β_1_-β_1_ interaction in CHOβ_1_ cells.

Then we asked whether ouabain has a direct or indirect effect on the β_1_-β_1_ interaction. Ouabain binding to the α subunit could induce a localized conformational change that would make the β_1_ subunit more adhesive; which then could be considered as a direct effect. For instance, fluorometry assays identified three positions on the β_1_ subunit, that indicate that α and β subunits move toward each other during conformational transition, and produce a conformal rearrangement [66, 67]. Additionally, ouabain binding produce conformational rearrangements in α subunit of the sodium pump [68]. To further support this idea, our *in vitro* experiments showed that although the α subunit, is present during the immobilization on Ni-NTA and pool down assays, ouabain treatment does not modify β_1_-β_1_ interaction. As such, our results suggest that ouabain does not modify the adhesion between β_1_ subunits directly and that it requires additional cellular components to fulfil this function. In this regard, binding of ouabain to α subunit could activate signaling cascades that would up-regulate the amount of Na^+^,K^+^-ATPase exposed to the extracellular space and thus increase indirectly cell-cell adhesion mediated by β_1_ subunits. During last years it was established that the Na^+^,K^+^-ATPase is not just a pump [31, 69, 70]. Experimental evidence strongly indicate that epithelial cells contain at least two populations of Na^+^,K^+^-ATPases: A pumping and a signaling (non-pumping) one [23,59,70]; which could also be the case for transfected CHOβ_1_ fibroblasts. Overexpression of β_1_ subunit up regulates the expression of α subunit [31, 71]. Both are assembled in endoplasmic reticulum and targeted to the plasma membrane [72]. Consistent with this idea, confocal microscopy and surface biotinylation analyses performed in this study show that ouabain stimulates the expression and delivery of the pump to the plasma membrane. Apparently, the signaling population is increased because we observed that even without ouabain the basal expression of cSrc, AKT, and ERK1/2 is increased in comparison to non-transfected CHO cells (Sup 1). Moreover, physiological concentrations of ouabain stimulated the activation of cSrc; furthermore, the pNaKtide which only binds to cSrc associated with Na^+^,K^+^-ATPase, successfully reverted the effect of ouabain on cSrc activation and cell-cell adhesion. In addition, AKT also seems to regulate the ouabain-dependent cell-cell adhesion observed in CHOβ_1_ fibroblasts, since the specific inhibitor of AKT abolished the β_1_-mediated cell-cell adhesion. This is not surprising as it has been shown that AKT trans-activates IP3K which forms part of the signaling complex in association with Na^+^,K^+^-ATPase [73,74]. On the other hand, the inhibition of ERK signaling cascade had a minor effect in the ouabain-dependent cellular aggregation. This might be partially explained by the fact that ERK1/2 is not the only signaling effector activated by cSrc [75,76]. Moreover, our initial screening revealed various signaling proteins that were not studied here and that are probably involved such as TrkB, EphB4 and PDGFR. Future studies will be directed to understand the participation of these signaling molecules in the β_1_-β_1_ adhesion phenotype.

All together our results suggest that overexpression of β_1_ subunit in CHO cells increases the population of Na^+^,K^+^-ATPases at the plasma membrane. This pool of pumps is most probably involved in mediating β_1_-β_1_ interactions between neighboring cells. Whether these proteins have an active role in ion transport remains to be elucidated. However, a plausible model would involve the activation of cSrc and AKT upon stimulation of CHOβ_1_ cells with physiological concentrations of ouabain. These signaling cascades would result in the overexpression of adherent pumps targeted to cell contacts at the plasma membrane. Nonetheless, an open question that would have to be addressed in the future is if CHOβ_1_ cells form signalosomes, as epithelial cells do and if β_1_-β_1_ interaction occurs between specific pools (pumping and non-pumping pool) or not.

## Materials and Methods

### Cell Culture

Starter Cell lines, MDCK (Madin-Darby Canine Kidney, CCL-34) and CHO-K1 (Chinese hamster ovary) were obtained from the American Type Culture Collection. CHO-K1 cells were cultured in a mixture of F12/DMEM (1132-082, Invitrogen), supplemented with 10% fetal calf serum (FCS) (200-6170, Invitrogen), 100 U/ml penicillin, and 100 μg/ml streptomycin (600-5145, Invitrogen) and MDCK cells were cultured in DMEM (430-1600 Invitrogen) supplemented with 10% fetal calf serum (FCS) (200-6170, Invitrogen), 100 U/ml penicillin, and 100 μg/ml streptomycin (600-5145, Invitrogen). Cells were grown at 36.5°C a 5% CO_2_ atmosphere, 90% humidity (Forma Scientific CO_2_ incubator, Steri-Cult 200).

For all experiments, the cells were depleted of serum 24 h before adding different concentrations of ouabain (10 nM, 50 nM, 100 nM 10 µM or 100 µM as indicated in the figures (3125, Sigma).

CHO-K1 cells were transfected with pCIN4-β_1_ (CHOβ_1_), (pIRESneo-His_6_ (CHOβ_1_ His_6_), and pEYFP-β_1_ dog (CHOβ_1_ YFP) plasmids as previously described [16, 46, 48], CHO–K1 cells transfected with EcDendra [49] were a kind gift of P. Nava Cinvestav, Mexico. The construct for secreted ecto-domain of dog β_1_ subunit (secβ_1_) was a kind gift of D. M. Fambrough, and was generated as previously reported [50]. Secβ_1_ was transfected in CHO cells as described in Padilla-Benavides 2010. CHO secβ_1_ cells were cultured in serum free media for 4 days before collecting the conditioned medium containing the ectodomain of β_1_ subunit (Secβ1) [16, 51]. Stable clones were selected and maintained with 0.2 mg/ml G418 (11811031, Invitro) in F12/DMEM mixture.

For experiments including secβ_1_ the CHOβ_1_ cells were seeded in 24 wells plates and after 90 minutes the monolayer was washed three times with PBS and a serum free medium with or without secβ_1_ was added. The next day the medium was replaced with a medium containing ouabain and secβ_1_.

The kinase inhibitors used were against cSrc (Dasatinib 100nM), AKT (Perifosine 10μM), ERK1/2 (U0126, 10μM) (0952, 14240 and 9903, respectively, Cell Signaling Technologies) and pNaKtide (1μM, kind gift of Dr. Z. Xie, Marshall Institute for Interdisciplinary Research). Inhibitors were added 1 h before ouabain treatment.

### Antibodies

The primary antibodies used were: mouse anti-α_1_-subunit of the Na^+^,K^+^-ATPase (7671, Abcam). Mouse anti-β_1_-subunit of the Na^+^,K^+^-ATPase (kind gift of Dr. M. Caplan, Yale University). Rabbit anti-phospho-Akt (Ser473, D9E), rabbit anti-phospho-p44/42 MAPK (Erk1/2, Thr202/Tyr204, D13.14.4E); rabbit anti-phospho-Src Family (Tyr416, D49G4); rabbit anti-Akt (pan, C67E7); rabbit anti-p44/42 MAPK (Erk1/2, 137F5), and rabbit anti-Src (36D10) (4060, 4370, 6943, 4691, 4695, and 2109, respectively, Cell Signaling Technologies).

The secondary antibodies used were: goat anti-rabbit IgG, and goat anti-mouse IgG coupled to HRP (65-6120 and 62-6520, respectively, Thermo Fisher Scientific). Goat anti-mouse IgG Alexa 488-conjugated (A11094, Molecular Probes).

### Immunofluorescence and confocal microscopy analyses

CHO and MDCK cells grown on coverslips at the indicated conditions in the figures, were immunostained as previously described [46]. Briefly, the cells were washed with PBS, fixed and permeabilized with ice-cold methanol for 5 min. After washing with PBS, the cells were blocked with 3% bovine serum albumin for 1 h, followed by 1 h incubation with a mouse primary antibody against the β_1_ subunit of the Na^+^,K^+^-ATPase (1:50 dilution) at room temperature. Cells were washed 10 times quickly with PBS, and incubated with a goat anti-mouse Alexa 488 secondary antibody (1:500 dilution) for 30 min at room temperature. After washing, the cells were mounted on glass slides with FluoroGuard antifade reagent (170–3140, BioRad). Samples were imaged using a confocal microscope (Leica TCS SP8), and visualized using the Leica Lite software. The relative fluorescence intensity was quantified using Fiji 1.0 software.

### Western blot analysis

Monolayers of CHO cells grown on multi-well plates were washed 3 times with ice-cold PBS and incubated in RIPA buffer (24948A, Santa Cruz) with protease inhibitors (2498, Santa Cruz) for 30 min under continuous and vigorous shaking. After that the cells were scraped and collected into a 1.5 ml microcentrifuge tube. The cell extract was homogenized with insulin syringe and centrifuged at 17,700 x g for 10 min. The supernatant was recovered, and protein concentration was quantified by BCA as indicated by the manufacturer (23225, Pierce Chemical). Thirty μg of protein were separated on 10% SDS-PAGE gels, electro-transferred onto a PVDF membrane (RPN 303F, Hybond-P Amersham Biosciences), and detected using the indicated primary (anti-α_1_-subunit of the Na^+^,K^+^-ATPase (1:2000), mouse anti-β_1_-subunit of the Na^+^,K^+^-ATPase (1:500). Rabbit anti-phospho-Akt (1:2000), rabbit anti-phospho-p44/42 MAPK (1:2000), rabbit anti-phospho-Src Family (1:2000), rabbit anti-Akt (1:2000), rabbit anti-p44/42 MAPK (1:2000), and rabbit anti-Src (1:2000) and appropriate secondary antibodies (goat anti-rabbit IgG, and goat anti-mouse IgG coupled to HRP (1:10000) and goat anti-mouse IgG Alexa 488-conjugated (1:10000). Membranes were developed with ECL PLUS (RPN2132, Amersham Biosciences). Immunoblots were imaged by FUSIONFX (Vilber), and ChemiDocXRS (BioRad) and quantified by densitometry using Fiji 1.0 software.

### Cell surface Biotinylation

Confluent monolayers of CHOβ_1_ cells were depleted of serum for 24 h and then incubated with or without 50 nM ouabain for 24 h. Then, the monolayers were washed 3 times with PBS, and incubated with 1 mg/ml of EZ-Link Sulfo-NHS-SS-Biotin (21331, Thermo scientific) for 30 min. Subsequently the monolayers were washed 3 times with PBS containing 100 mM glycine to quench the excess of Biotin, followed by a final wash with PBS containing 1% Triton X-100 and protease inhibitors. After 30 min, the cells were scraped, and the cell lysate collected into a 1.5 ml microcentrifuge tubes. The extract was homogenized by passing it 10 times through insulin syringe and centrifuged for 10 min at 9,000 x g at 4 °C. The supernatant was recovered, and the protein content was measured using the BCA protein assay method. The biotinylated extracts were incubated overnight at 4 °C with 100 μl of streptavidin-agarose suspension (S1638, Sigma). The next day, bead-adherent complexes were washed 5 times with PBS, and finally the proteins were eluted in 2X Laemmli buffer and boiling for 5 min.

### Cell adhesion assay (Dispase assay)

Confluent Monolayers of CHOβ_1_ cells seeded on 24 well plates were depleted of serum for 24 h, and then incubated with or without ouabain (50nM) for 24 h. Then the monolayers were washed with ice-cold PBS, and detached from the plates by incubation with PBS without Ca^2+^ supplemented with 0.6 U/ml of Dispase I (D4818, Sigma) for 35 min at 37 °C. Subsequently the Dispase solution was carefully removed using a 200 µl pipette tip, and replaced by 100 μl of PBS. The cells were then mechanically stressed by pipetting up and down 5 times using a 200 µl pipette. The resulting aggregates were visualized by light microscopy using the 10 X and 20 X objectives (Axiovert 200M Fluorescence/Live Cell Imaging, Carl Zeiss). Three independent biological replicates were imaged using the AxioVision 4.8 sofware. The number of aggregates was counted using the Fiji 1.0 software cell counter (aggregates <200μm^2^ were excluded from the quantification).

### Pull Down assay (PD)

Total extract of CHO cells expressing the β_1_His_6_ construct were immobilized with nickel-nitrilotriacetic acid beads (Ni^+^-NTA, His Trap FF column; GE Healthcare) equilibrated with 10 ml of RIPA buffer containing protease inhibitors. Total protein extracts (500 µg) were loaded into the resin, and allowed to interact for 3 h at 4°C with gentle shaking. Then the unbound protein was washed as indicated by the manufacturer, and the total extract of CHOβ_1_ YFP cells incubated in the presence or absence of ouabain (50 nM and 100 µM) were loaded as a prey. Samples were incubated overnight at 4°C, and washed with PBS supplemented with 10 and 20 mM imidazole. Interacting proteins were eluted in PBS containing 500 mM imidazole and were analyzed by western blot.

### Statistical analysis

GraphPad Prism version 7.00 software was used for all statistical analyses. The data are presented, as the mean ± SEM or SD as indicated in the figures. Statistical significance was determined using ANOVA one way and Kruskal Wallis test, and *t*-test for two conditions. *P* ≤ 0.05 was considered significant.

## Supporting information

Vilchis-Nestor Supplemental Information

## Author Contributions

Conceptualization, L.S. and C.A.V-N.; methodology, CA V-N.; investigation, C.A. V-N. and M.L. R.; resources, L.S. and T P-B.; writing—original draft preparation, L.S. and CA V-N.; writing—review and editing, and T.P-B.; supervision, L.S and T.P-B.; project administration, L.S..; funding acquisition, L.S and T.P-B.

## Acknowledgments

The authors thank Xie Z. for the pNaKtide and Rosalia Aguirre for her dedicated technical assistance.

