## Supplementary material for "Ouabain enhances cell-cell adhesion mediated by β_1_-subunits of the Na^+^,K^+^-ATPase in CHO fibroblasts": Vilchis-Nestor Supplemental Information

### FIGURE S1

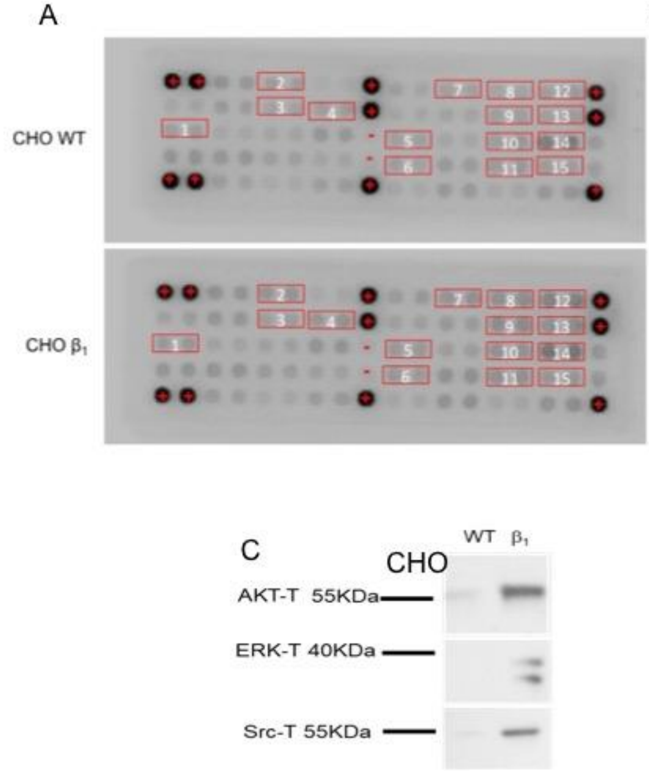


B

| | Target | Phosphorilation site | Family |
|---|---|---|---|
| 1 | HER2/ErB2 | Pan-Tyr | EGFR |
| 2 | TrkA/NTRK1 | Pan-Tyr | NGFR |
| 3 | TrkB/NTRK2 | Pan-Tyr | NGFR |
| 4 | PDGFR | Pan-Tyr | PDGFR |
| 5 | EphA3 | Pan-Tyr | EphR |
| 6 | EphB1 | Pan-Tyr | EphR |
| 7 | EphB4 | Pan-Tyr | EPhR |
| 8 | AKT/PKB/Rac | Thr 308 | Akt |
| 9 | AKT/PKB/Rac | Ser 473 | Akt |
| 10 | p44742 MAPK(ERK1/2) | Thr 202/tyr 204 | MAPK |
| 11 | S6 Ribosomal protein | Ser 335/236 | RSK |
| 12 | IRS-1 | Pan-Tyr | IRS |
| 13 | Zap-70 | Pan-Tyr | Zap-70 |
| 14 | Src | Pan-Tyr | Src |
| 15 | Lck | Pan-Tyr | Src |

**Figure A1.** Phosphorylation state of major Receptor Tyrosine Kinases (RTKs) in CHO WT, and CHOβ$_1$ cells. A) The RTK signaling antibody array performed with total extracts of CHO WT and CHOβ$_1$ cells incubated for 24 h without serum. Each point represents one RTK (PathScan RTK Signaling Antibody Array Kit) B) List of proteins that apparently increase their phosphorylated state in CHOβ$_1$ cells compared to CHO WT cells. C) Representative Western blot (n=2) showing the expression of AKT-T, ERK 1/2-T and Src-T in CHO WT and CHOβ$_1$ cells with specific antibodies.

### FIGURE S2

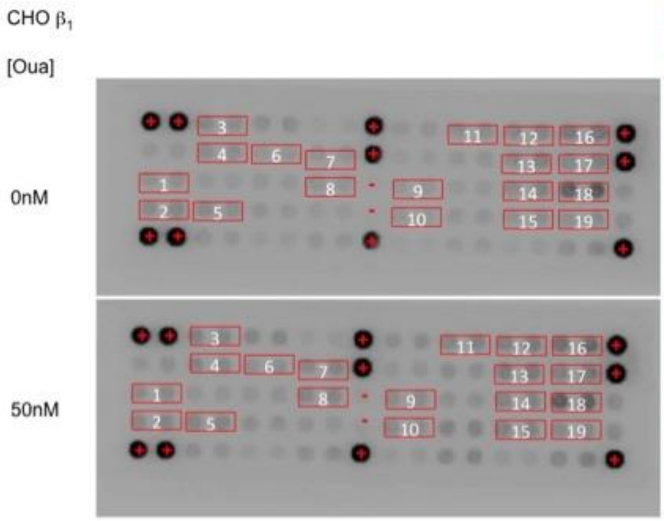


| | Target x | Phosphorilation site | Family |
|---|---|---|---|
| 1 | HER2/ErB2 | Pan-Tyr | EGFR |
| 2 | HER3/ErbB3 | Pan-Tyr | EGFR |
| 3 | FGFR1 | Pan-Tyr | FGFR |
| 4 | FGFR3 | Pan-Tyr | FGFR |
| 5 | InsR | Pan-Tyr | Insulin R |
| 6 | Trkb/NTRK2 | Pan-Tyr | NGFR |
| 7 | PDGFR | Pan-Tyr | PDGFR |
| 8 | c-Kit/SCFR | Pan-Tyr | PDGFR |
| 9 | EphA3 | Pan-Tyr | EphR |
| 10 | EphB1 | Pan-Tyr | EphR |
| 11 | EphB4 | Pan-Tyr | EPhR |
| 12 | AKT/PKB/Rac | Thr 308 | Akt |
| 13 | AKT/PKB/Rac | Ser 473 | Akt |
| 14 | p44742 MAPK(ERK1/2) | Thr 202/tyr 204 | MAPK |
| 15 | S6 Ribosomal protein | Ser 335/236 | RSK |
| 16 | IRS-1 | Pan-Tyr | IRS |
| 17 | Zap-70 | Pan-Tyr | Zap-70 |
| 18 | Src | Pan-Tyr | Src |
| 19 | Lck | Pan-Tyr | Src |

**Figure A2.** Phosphorylation status of major RTKs in CHOβ1 cells treated or not with ouabain. A) The RTK Signaling Antibody Arrangement was performed with total extracts of CHO β1 cells treated with and without 50 nM ouabain for 24 hr. B) List of proteins that apparently modified their phosphorylated state in CHO β1 cells treated with 50 nM of ouabain compared to CHO β1 cells without ouabain.

#### Supplemental methods

##### Phospho-RTK Antibody Array

A PathScan RTK Signaling Antibody Array Kit (7982, Cell Signaling Technology) was used according to the manufacturer’s instructions to assess the phosphorylation status of the major RTKs. Briefly, cell lysate was incubated on a slide, to which was added a biotinylated detection antibody mixture, streptavidin-conjugated horseradish peroxidase, and, finally, enhanced chemiluminescence reagents. Slide images were captured with ChemiDocXRS (BioRad).
